# *hTERT* Expression, Regulation, and Prognostic Significance in Pediatric Medulloblastoma

**DOI:** 10.64898/2026.05.01.722294

**Authors:** Ryuma Tanaka, Banlanjo Umaru, Matthew Sobo, Shiva Senthil Kumar, Kathleen Dorris, Volker Hovestadt, Vijay Ramaswamy, Marc Remke, Ashley Margol, Charles B. Stevenson, Shahab Asgharzadeh, Stewart Goldman, Lili Miles, Jie Huang, Katja von Hoff, Stefan Rutkowski, Arzu Onar-Thomas, Uri Tabori, Michael Taylor, Stefan M. Pfister, Ralph Salloum, Maryam Fouladi, Rachid Drissi

**Author notes:** Corresponding author **Correspondence to:** Rachid Drissi, PhD, Associate Professor, Center for Childhood Cancer Research, Nationwide Children’s Hospital, 575 Children’s Drive, Columbus Ohio. These authors contributed equally (Ryuma Tanaka, Banlanjo Umaru).

## Abstract

**Background:** Telomerase reactivation, a hallmark of many cancers, is associated with expression of its catalytic subunit, hTERT. However, the prognostic significance of telomere maintenance mechanisms in pediatric medulloblastoma remains poorly defined.

**Methods:** In this multi-institutional retrospective study of telomerase expression and *hTERT* regulation in newly diagnosed children with medulloblastoma, *hTERT* and *MYC* expression were assessed by qRT-PCR, normalized to non-neoplastic brain control samples. *hTERT* promoter methylation was analyzed using quantitative pyrosequencing and Illumina 450k methylation array. Cox proportional-hazard regression analyses evaluated the association of *hTERT* expression with progression-free survival (PFS) or overall survival (OS). Spearman correlation and Kruskal-Wallis tests correlated *hTERT* promoter methylation and expression and assessed variations among medulloblastoma subgroups, respectively.

**Results:** Among 74 patients with available *hTERT* expression and outcome data, higher expression was associated with worse OS (HR=1.22, 95% CI: 1.01-1.47, p=0.036) and PFS (HR=1.17, 95% CI: 1.00-1.37, p=0.051) after adjusting for subgroup. Similar results were obtained when adjusting for metastatic status. Group 3 patients had the highest *hTERT* expression (p=0.001). Pyrosequencing data were available for 61 patients and 450k methylation array data for 292 patients. *hTERT* promoter was differentially methylated across subgroups with WNT followed by group 3 demonstrating the highest methylation on 450k (p<0.0001), findings that were confirmed by pyrosequencing. *hTERT* promoter methylation positively correlated with *hTERT* expression (Spearman correlation=0.42, p=0.02 by 450k and 0.34, p=0.007 by pyrosequencing). No significant correlation was observed between *hTERT* and *MYC* expression.

**Conclusion:** Elevated *hTERT* expression is associated with worse PFS and OS in medulloblastoma across subgroups, supporting telomerase inhibition as a potential therapeutic strategy.

## BACKGROUND

Medulloblastoma is an embryonal tumor of the cerebellum that accounts for around 20% of brain tumors in children. The current management of medulloblastoma is based on traditional prognostic factors used to risk stratify patients: age, extent of resection, presence of metastatic disease and histology[13, 45]. Patients with standard-risk disease have a 5-year event free survivals (EFS) of 79%-85%[25-27], whereas high-risk patients have a 5-year EFS ranging between 55% to 70% at best despite intensive multimodal therapy[15, 45] . Seminal work by Pomeroy et al.[30], Cho et al.[7], and Kool et al.[21] has elucidated the genomic landscape of medulloblastoma leading to a consensus recognition of 4 subgroups (WNT, SHH, group 3 and group 4) with distinct gene expression profiles, methylomes and clinicopathological features[41]. Based on these findings, risk classifications that incorporate biological subgrouping as well clinicoradiological factors are being adopted for stratification of patients with medulloblastoma. Despite these advances, identification of actionable genomic aberrations and the integration of novel molecularly targeted agents into therapeutic regimens have remained limited.

One such potential target is telomerase, a reverse transcriptase complex that includes a catalytic subunit encoded by the *hTERT* gene and an RNA template (TERC)[4]. Telomerase replenishes shortening telomeres by restoring DNA at the end of chromosomes with each cell division, thereby preventing cells from undergoing senescence[9]. Telomerase activity is highly correlated with *hTERT* mRNA expression[40, 44] and is almost universally detected in cancer whereas normal somatic tissue, including brain, does not demonstrate telomerase activity[3, 5, 19, 38]. In children with central nervous system (CNS) malignancies, high telomerase expression has been shown to be a poor prognostic marker in many tumor types. Specifically, initial studies in pediatric intracranial ependymomas revealed *hTERT* expression as the single most important predictor of outcome in multivariate analysis[39]. A more recent analysis accounting for molecular subgroups of ependymoma by Gojo et al.[16] showed a significant association between increased telomerase activity and dismal prognosis in patients with posterior fossa group A ependymoma. We have previously established that telomerase activity is associated with shorter overall survival (OS) in children with high-grade gliomas after controlling for tumor grade and extent of resection[12]. However, few studies have evaluated telomerase activity in pediatric medulloblastoma. In one study, upregulation of *hTERT* mRNA was demonstrated in the majority (76%) of primitive neuroectodermal tumors (n=50) including 43 medulloblastoma samples[10]. Fan et al.[14] showed a trend towards shorter survival in medulloblastoma patients (n=38) with high telomerase expression, though the relatively short median follow-up of 20 months is a potential limitation. Many mechanisms have been implicated in the upregulation of telomerase activity including *hTERT* promoter methylation[3, 6] and mutation[18], *hTERT* amplification[46] and genomic rearrangement[28]. In this multi-institutional study, we present findings on telomere maintenance mechanisms and *hTERT* expression, and their association with clinical outcomes, in a large cohort of children with newly-diagnosed medulloblastoma.

## MATERIALS AND METHODS

### Study cohort

We conducted a retrospective study of 79 children with medulloblastoma diagnosed between 1990 and 2011 who had available tumor tissue at Cincinnati Children’s Hospital Medical Center (CCHMC; Cincinnati, OH), Ann and Robert H. Lurie Children’s Hospital of Chicago (Chicago, IL), Children’s Hospital of Los Angeles (CHLA; Los Angeles, CA), or Brain Tumor Tissue Bank (Ontario, Canada). Institutional review board approval was obtained at CCHMC and at the participating institutions as applicable. Data generated using samples at these institutions was supplemented by additional genomic data from 340 children with medulloblastoma provided by the German Cancer Research Centers in Heidelberg and Düsseldorf, Germany, the Labatt’s Brain Tumour Research Centre at the Hospital for Sick Children, Toronto, Canada. All specimens were collected in accordance with Cincinnati Children’s Hospital Medical Center Institutional review board (Study ID: 2010-1597). Patient identifiers were removed prior to evaluation, and a single sequential numerical identifier was assigned to each patient. Institutional review board approval was obtained at CCHMC and at the participating institutions. All tumors were centrally reviewed by two experienced neuropathologists to confirm the diagnosis of medulloblastoma.

### *hTERT* promoter mutation and methylation

Extraction of DNA was performed as previously described[12]. Region of the *hTERT* promoter harboring the recurrent mutations C228T and C250T were (Forward: 5’-AGCACCTCGCGGTAGTGG-3’, Reverse: 5’-GTCCTGCCCCTTCACCTT-3’). Sanger sequencing was performed using the amplified products. *hTERT* promoter methylation status was performed using quantitative pyrosequencing and β-values derived from the 450K data.

### Medulloblastoma subgrouping 450K methylation profile and other methods

Whole genome methylation was carried out using Illumina HumanMethylation450K (Illumina) by the Sequencing and Genotyping Core (Cincinnati Children’s Hospital). Raw values were preprocessed by the stratified quantile normalization function in the minfi package (Bioconductor) in the R environment. Patients from the CHLA cohort were placed into subgroups using Taqman Low Density Array. Other patients were sub-grouped based on the 450k (Illumina) methylation status 48-probe signature described by Hovestadt[17]. WNT and SHH patients were confirmed using immunohistochemistry directed at β-Catenin and Gab1 respectively.

### *hTERT, c-MYC*, Expression

Total RNA was extracted by RNAzol reagent (Molecular Research Center, Cincinnati, OH), and one μg RNA was converted to cDNA using First Strand cDNA Synthesis kit (USB, Cleveland, Ohio). mRNA expression analysis was performed by qRT-PCR using primer/probe sets that spans exon junctions (*hTERT*: Hs00972650_m1, *c-MYC*: Hs00153408_m1, Applied Biosystems, Carlsbad, CA). GAPDH (Hs03929097_g1 Applied Biosystems, Carlsbad, CA) was used as the housekeeping gene. qRT-PCR assays were performed for each sample at least twice in triplicate. Relative expression (expressed as RQ) levels of *hTERT* mRNA and *cMYC* mRNA were compared to the mean expression level from non-neoplastic control brain samples using the ΔΔCt method[35]. RQ values >1 defined increased expression levels as compared to the mean values for control non-neoplastic specimens.

### TRAP Assay

Telomerase enzyme activity was assessed in MB tumor tissue samples by Telomeric Repeat Amplification Protocol (TRAP) assay using the TRAPeze-XL Telomerase Detection Kit (Millipore, Cat. #S7707, Billerica, MA) as previously described [12]. HeLa cell line (30 ng total protein) and CHAPS buffer were used as positive and negative controls, respectively. Telomerase enzyme activity quantification was performed using the fluorometric detection following the user manual. Briefly, a standard curve was generated using measurements from the TSR8 template dilutions together with controls. The Total Product Generated (TPG) values for each sample were calculated from the standard curve. A unit of TPG corresponds to the number of template substrate extended with at least three telomeric repeats by the telomerase enzyme.

### Statistical Analyses

Cox proportional hazard regression analyses evaluated the association between the variables (*hTERT* expression, *hTERT* promoter methylation, *c-MYC* expression, and *c-MYC* amplification and *c-MYC* expression levels) with progression-free survival (PFS) or overall survival (OS). Because *hTERT* expression was highly right-skewed, values were log-transformed prior to analysis to reduce the influence of extreme outliers. PFS was defined as time from diagnosis to first disease progression or last follow-up visit without progression. OS was defined as time from diagnosis to death from disease or last follow-up visit. In patients without progression or death, PFS and OS were treated as right-censored, respectively.

Kruskal-Wallis test was used for assessing the variation of *hTERT* expression and *hTERT* promoter methylation status among MB subgroups. Spearman correlation was used to assess correlation between *hTERT* promoter methylation and *hTERT* expression; *hTERT* expression and *c-MYC* expression *hTERT* expression and *c-MYC* amplification. Results with p values <0.05 were reported as statistically significant.

## RESULTS

### *hTERT* is differentially expressed across medulloblastoma subgroups and is associated with poor prognosis

Among 78 patients with available *hTERT* expression, molecular subgroup classification (WNT, SHH, Group 3, Group 4), and clinical outcome data, 74 (95%) demonstrated detectable *hTERT* expression. Expression levels varied significantly across subgroups, with Group 3 tumors exhibiting the highest overall expression compared to other subgroups (Figure 1A; Kruskal–Wallis p=0.001). In a subset of samples with available material (n = 27), telomerase activity was assessed using the TRAP assay and based on analysis of the total product generated (TPG), telomerase enzyme activity was detected in 25/27 (92.6%) of the cases (Supplemental Figure 1). This data aligns with prior studies demonstrating strong correlation of telomerase activity with *hTERT* mRNA expression[40, 44]. The TRAP assay is highly sensitive; however, its reliability may be compromised in inadequately processed or preserved tissue samples due to denaturation or degradation of telomerase components. Cox proportional hazards regression models, incorporating log-transformed *hTERT* expression as a continuous variable and including subgroup as a categorical covariate, showed that higher *hTERT* expression was significantly associated with worse overall survival (OS; HR = 1.22, 95% CI: 1.01–1.47, p = 0.036), and showed a marginal association with worse progression-free survival (PFS; HR = 1.17, 95% CI: 1.00–1.37, p = 0.051), independent of subgroup affiliation. Kaplan–Meier analyses further illustrated these relationships, with patients dichotomized into high and low expression groups based on the median expression level as the cutoff (Figure 1B–C). Given the well-established prognostic impact of metastatic disease in medulloblastoma, additional models adjusting for metastatic status were performed. In this analysis, elevated *hTERT* expression remained independently associated with significantly worse OS (HR=1.35, 95% CI: 1.12-1.62, p=0.002) and PFS (HR=1.25, 95% CI: 1.07-1.45, p=0.005). Kaplan–Meier survival curves stratified by *hTERT* expression groups and metastatic status are shown in Supplemental Figure 2. Collectively, these findings support *hTERT* expression as a potential biomarker of aggressive disease in medulloblastoma.

**Figure 1.**
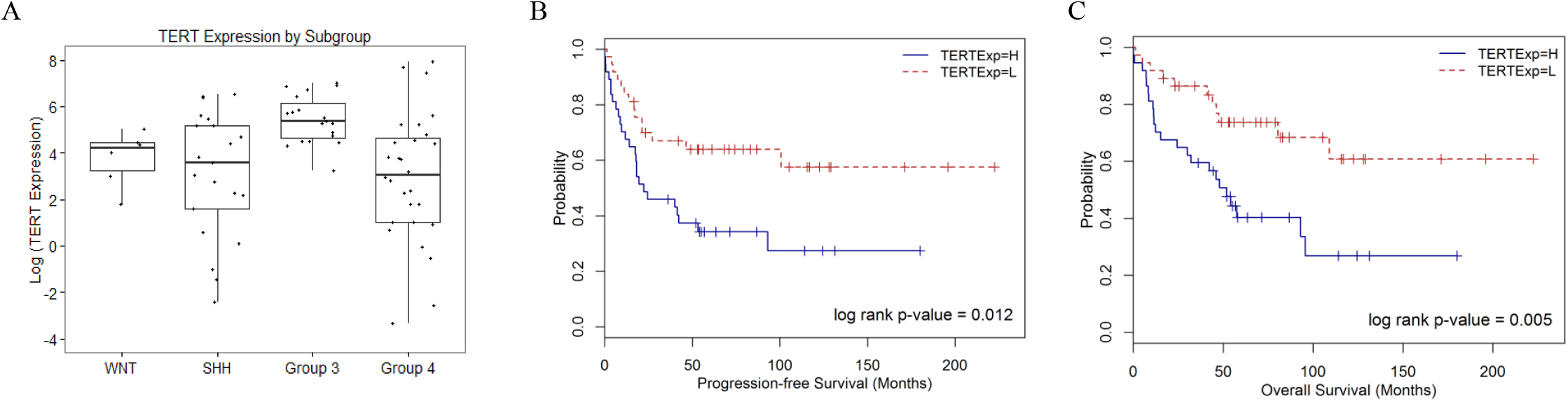
*hTERT* expression across medulloblastoma subgroups and its association with OS and PFS. All patients with available subgrouping, survival, and *hTERT* expression data were included in this analysis (n=74). (A) Gene expression analysis of *hTERT* in different medulloblastoma subgroups. Group 3 patients had higher *hTERT* expression compared with patients in the other 3 subgroups (Kruskal-Wallis test p-value = 0.001). (B-C) Overall survival (OS) and progression-free survival (PFS) by Kaplan-Meier analysis using log-rank statistics stratified by *TERT* expression. Patients were divided into high *TERT* expression (TERTExp=H) and low *hTERT* expression (TERTExp=L) groups using median expression as cutoff. Higher *TERT* expression was associated with worse OS (p=0.005) and PFS (p=0.012).

### *hTERT* Promoter Methylation Status

Given prior reports demonstrating the prognostic relevance of *hTERT* promoter hypermethylation in pediatric brain tumors[6, 16], we evaluated hTERT promoter methylation in patients with available 450k methylation (n=292) and pyrosequencing data (n=61) as described in the Methods. Analysis of the 450k dataset revealed significant differences in *hTERT* promoter methylation across molecular subgroups, with the highest methylation levels observed in WNT tumors, followed by Group 3 (Figure 2A, Kruskal-Wallis test p-value < 0.0001). These findings indicate that the *hTERT* promoter methylation is differentially methylated among subgroups. Pyrosequencing data confirmed these trends (Figure 2B). As expected, non-neoplastic brain specimens (n=6) did not exhibit increased methylation at the interrogated CpG methylation sites (data not shown). Among patients with available 450k methylation and survival data (n=200), *hTERT* promoter methylation status was not associated with OS or PFS across subgroups when analyzed as a continuous variable using Cox proportional hazards regression. Similarly, within each subgroup, dichotomized methylation status (hypermethylated vs hypomethylated) was not significantly associated with OS or PFS (Figure 3A–B). On multivariate analysis adjusting for subgroup, metastatic status and patient age, *hTERT* promoter methylation status did not carry any prognostic significance. Consistent results were observed when adjusting for metastatic status alone, with no association between dichotomized methylation status and survival outcomes (Figure 3C). These findings were corroborated using pyrosequencing data (Supplemental Figure 3). We next evaluated the relationship between *hTERT* promoter methylation and *hTERT* mRNA expression by qRT-PCR. Paired methylation and expression data were available for 30 patients (450k array) and 61 patients (pyrosequencing). *hTERT* promoter methylation demonstrated a moderate positive correlation with *hTERT* expression (Spearman correlation=0.42, p=0.02 by 450k and 0.34, p=0.007 by pyrosequencing) (Figure 4).

**Figure 2.**
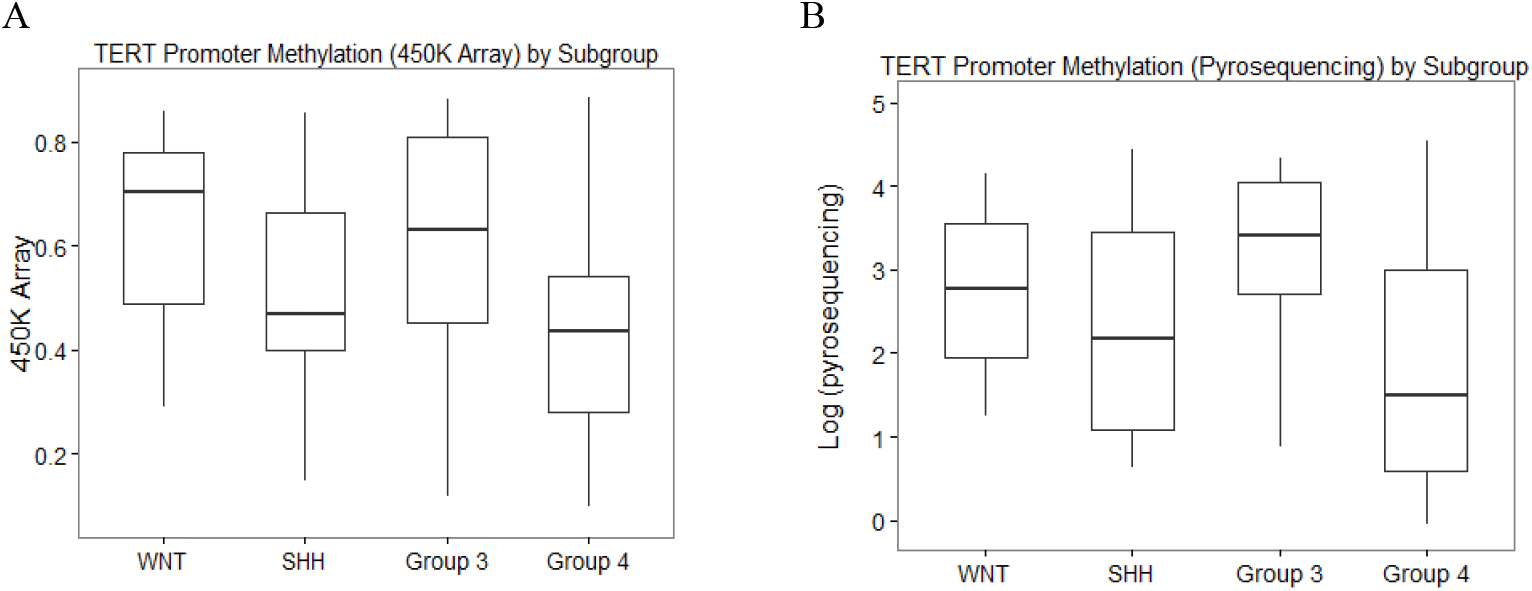
*hTERT* promoter methylation status among medulloblastoma subgroups. All patients with available subgroup and *hTERT* promoter methylation data were included in this analysis. TERT promoter methylation status was determined using the 450K array (n=292) (A) and pyrosequencing (n=60) (B). (A) WNT patients have the highest median value of *hTERT* promoter methylation by 450K, followed by patients in Group 3 (Kruskal-Wallis test p-value < 0.0001). (B) *hTERT* promoter as measured by pyrosequencing appears to be differentially methylated in medulloblastoma subgroups (Kruskal-Wallis test p-value = 0.055). Group 3 patients have the highest median value of *hTERT* promoter methylation.

**Figure 3.**
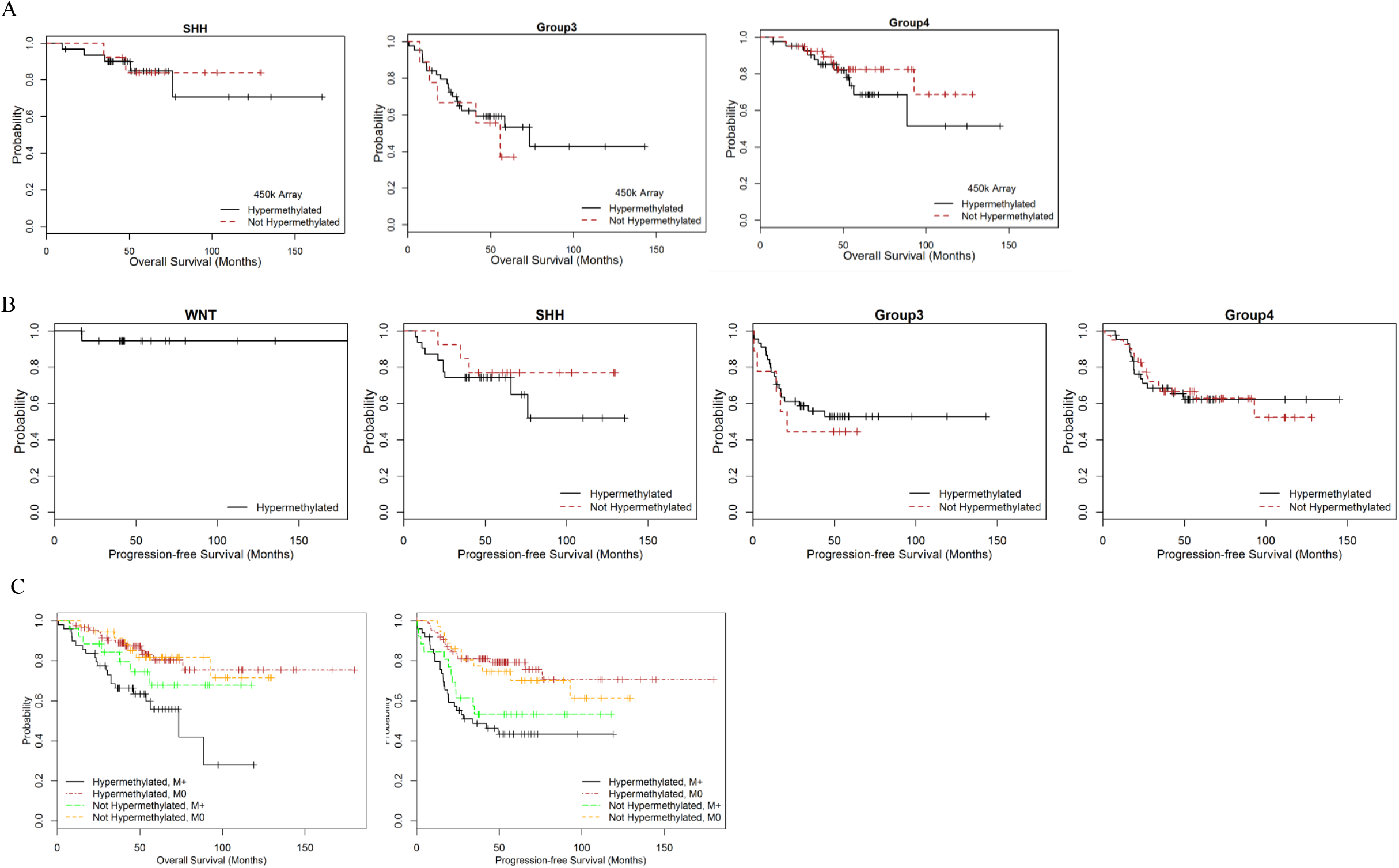
Association between *hTERT* promoter methylation status and survival outcomes. All patients with well-defined subgroup (WNT, SHH, Group 3, and Group 4), non-missing OS or PFS data, and non-missing 450k array data, are included in this analysis (n = 200, 49 OS events, 69 PFS events). (A-B) Kaplan-Meier plot showing the OS distribution (A) and PFS distribution (B) stratified by the *hTERT* promoter methylation status (hypermethylation and not hypermethylated) in different medulloblastoma subgroups. (C) Kaplan-Meier plot OS and PFS adjusted for metastatic status and *hTERT* promoter methylation status.

**Figure 4.**
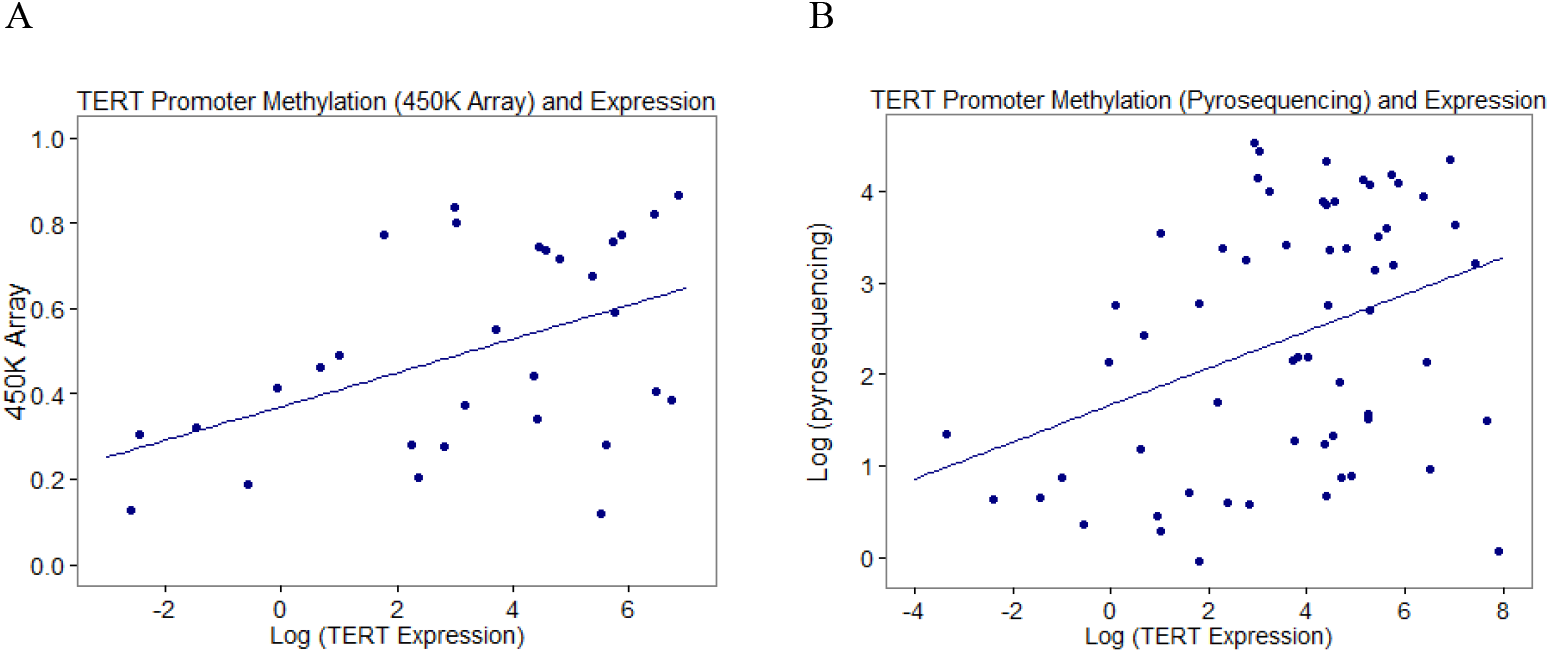
Correlation between *hTERT* promoter methylation and *hTERT* expression. All patients with available *hTERT* expression and *hTERT* promoter methylation data were included in this analysis. TERT promoter methylation status was determined using the 450K array (n=30) and pyrosequencing (n=61). (A-B) Scatter plots showing a moderate positive correlation between *hTERT* expression and *hTERT* promoter methylation levels determined by (A) 450K array (Spearman’s correlation ρ = 0.42, p = 0.020), and (B) pyrosequencing (Spearman’s correlation ρ = 0.34, p = 0.007).

### Other mechanisms regulating *hTERT* expression

Hotspot mutations in *hTERT* promoter (C228T/A and C250T) have been reported in many malignancies[42] and are recurrent in adult SHH medulloblastoma[20, 22, 31]. In the present study, among a subset of samples (n=78, ages: 4 months-16 years) with available *hTERT* promoter sequencing data, only 2 tumors (2.6%) were found to harbor *hTERT* promoter mutations (C228Tmutation in both cases). Consistent with previous studies, both tumors belonged to the SHH subgroup, did not exhibit *hTERT* promoter hypermethylation and demonstrated elevated *hTERT* expression.

Alterations in *MYC* and its paralog *MYCN* have been well described in medulloblastoma[24, 29]. *MYC*, a marker of aggressive disease in Group 3 medulloblastoma[7], has also been implicated as a transcriptional regulator of *hTERT* in many cancer types[2, 33]. We therefore evaluated the association between *MYC* expression, *MYC* amplification and *hTERT* expression. Among 74 patients with available data, no significant correlation was observed between *MYC* and *hTERT* expression (Spearman correlation=0.15, p-value = 0.205) (Supplemental Figure 4A). Similarly, no association was identified between *MYC* amplification and *hTERT* expression in the subset of patients with available genomic data (n=34; Spearman correlation= -0.009, p-value = 0.959) (Supplemental Figure 4B).

To evaluate potential interactions among multiple mechanisms regulating *hTERT* expression, the 48 patients tested for *MYC* expression, *MYC* amplification, *hTERT* promoter mutation, and *hTERT* promoter methylation were stratified according to *hTERT* mRNA expression levels into five categories: Baseline, Low (<10× baseline), Moderate (10–50× baseline), High (50–100× baseline), and Very high (>100× baseline), (Figure 5). Baseline expressers (n = 4): Three of the four patients harbored an alteration in a single regulatory pathway (elevated *MYC* expression), while one patient had no identifiable abnormalities across the four assessed regulatory mechanisms. Low expressers (n = 10): Eight patients demonstrated a single altered mechanism (three with *hTERT* promoter hypermethylation and five with elevated *MYC* expression). One tumor was linked to both mechanisms, and one could not be linked to any of the regulatory pathways tested. Moderate expressers (n = 18): Four tumors could not be associated with known regulatory mechanisms. Seven patients showed a single altered mechanism (*hTERT* promoter hypermethylation or elevated *MYC* expression). Six tumors demonstrated two mechanisms, elevated *MYC* expression together with either *hTERT* promoter hypermethylation (n = 5) or *MYC* amplification (n = 1). One tumor exhibited all three abnormalities. High expressers (n = 10): One patient lacked identifiable regulatory alterations. Two patients had a single altered pathway (*hTERT* promoter hypermethylation or *MYC* expression). Five patients demonstrated two altered mechanisms (*hTERT* promoter hypermethylation with *MYC* expression or *hTERT* promoter mutation, or *MYC* expression with *MYC* amplification). Two patients had alterations in three regulatory systems (*hTERT* promoter hypermethylation with elevated *MYC* expression and *MYC* amplification). Very high expressers (n = 6): One patient showed no changes in any of the mechanisms evaluated. Two patients demonstrated a single altered mechanism (*MYC* expression or *hTERT* promoter mutation). Three patients displayed alterations in two mechanisms (e.g., *MYC* expression with *hTERT* promoter mutation or *hTERT* promoter hypermethylation). Importantly, *hTERT* promoter mutations were consistently associated with high or very high *hTERT* expression, regardless of other regulatory abnormalities. However, the presence of two patients with high or very high *hTERT* expression without any detectable regulatory alteration suggests that additional, suggests that additional, uncharacterized pathways likely contribute to *hTERT* activation.

**Figure 5.**
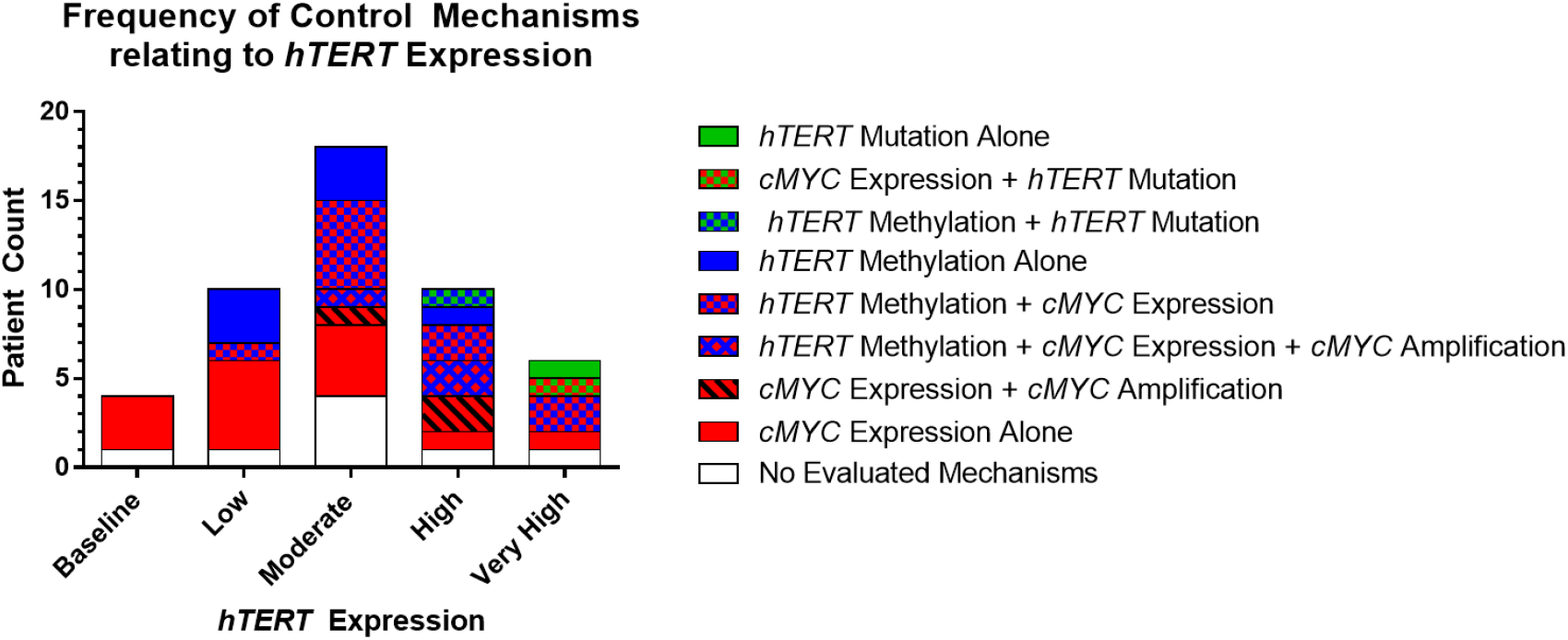
Distribution of putative regulatory mechanisms associated with *hTERT* expression levels. Stacked bar chart illustrates the distribution of genetic and epigenetic alterations associated with *hTERT* expression levels. Patients were stratified into five categories based on *hTERT* transcript abundance (Baseline, Low, Moderate, High, Very High). Bars indicate the number of patients per group, stratified by *hTERT* mutation, promoter methylation, *cMYC* expression and/or amplification, their combinations, or absence of identifiable mechanisms. Higher *hTERT* expression levels are enriched for combinatorial regulatory alterations, whereas baseline and low expression groups are predominantly characterized by isolated *c-MYC* expression or no detectable mechanism.

## DISCUSSION

This study is the first to characterize *hTERT* expression and its prognostic significance in a large cohort of children with newly diagnosed medulloblastoma. Our findings demonstrate that increased *hTERT* mRNA expression is associated with significantly worse OS and PFS, independent of molecular subgroup. These results are consistent with observations in other pediatric malignancies, including neuroblastoma and Wilms’ tumor[8, 11]. In contrast, Gojo et al. recently reported that in children with intracranial ependymoma only telomerase activity as measured by the telomerase repeat amplifying protocol assay was associated with survival, whereas *hTERT* mRNA levels assessed by qRT-PCR was not[16]. Given the well-established correlation between *hTERT* expression and telomerase activity[3, 44], this discrepancy may be attributable to the small sample size in the ependymomas cohort analyzed (n=22). Notably, in that study, the five patients lacking detectable *hTERT* mRNA also exhibited no telomerase activity and remained progression-free at 10-year follow-up.

The prognostic significance of telomerase activity and *hTERT* mRNA expression likely reflects underlying aggressive tumor biology, consistent with the established role of telomerase reactivation in oncogenesis. In addition, telomerase activity may contribute to treatment resistance, as it has been associated with reduced tumor sensitivity to both radiation and chemotherapy[43, 47]. Castelo-Branco et al.[6] showed that hypermethylation status of the *hTERT* promoter at five CpG sites upstream of the transcription start site (UTSS) is specific to malignant tumors and predicts poor prognosis in pediatric brain tumors. Among these, the CpG site cg11625005 was shown to closely represent the methylation status of the surrounding promoter region. However, the relationship between *hTERT* promoter hypermethylation and gene expression remains inconsistent across studies. while a positive association was reported by Castelo-Branco et al., other investigations in pediatric and adult brain tumors have described absent or even inverse correlations.[1, 16]. In our study, the methylation status at the key cg11625005 CpG site was positively correlated with *hTERT* expression, consistent with Lindsey et al’s findings in medulloblastoma [22]. Despite this, *hTERT* promoter hypermethylation did not demonstrate prognostic significance. This likely reflects the complex and multilayered regulation of *hTERT* transcription, of which promoter methylation represents only one component. In contrast, *hTERT* expression may better capture the integrated effects of epigenetic modification, transcriptional regulation, and tumor heterogeneity, and thus more accurately reflect biologically relevant telomerase activity.

The low frequency of *hTERT* promoter mutations observed in our cohort (2/78, 2.6% of all tested tumors and 2/19, 10.5% of SHH tumors) is consistent with prior reports demonstrating that these alterations are largely restricted to the SHH subgroup and are enriched in adult patients[20, 22, 31]. The rarity of *hTERT* promoter mutations in pediatric medulloblastoma, together with the elevated levels of *hTERT* promoter methylation observed across subgroups, underscores the prominent role of epigenetic dysregulation in the pathogenesis of pediatric brain tumors. Mechanistically, *hTERT* promoter methylation has been paradoxically associated with increased gene expression, potentially through inhibition of CTCF binding, a transcriptional repressor, to methylated DNA [32].

This study is limited by the lack of uniformity in treatment across patient cohorts. However, this variability is partially mitigated by the overall similarity of medulloblastoma treatment protocols across Europe and North America. Additionally, other regulatory mechanisms, such as *hTERT* promoter rearrangements, were not assessed and may contribute to telomerase activation.

In conclusion, our findings support telomerase inhibition or telomerase-dependent telomere targeting as potential therapeutic strategies across medulloblastoma subgroups. Although our prior work demonstrated that Imetelstat (Geron, Menlo Park, CA), a telomerase inhibitor targeting the template region of TERC, achieved intratumoral target engagement, its clinical utility in children with recurrent or refractory CNS tumors was limited by toxicity[34, 36]. Alternative approaches to targeting telomerase remain of interest. For example, induction of telomere dysfunction using 6-thio-deoxyguanosine, a guanine analog that functions as a telomerase substrate precursor[23], was tested in preclinical models of pediatric brain tumors[37].

## Acknowledgements

We thank the children and families who have generously donated tumor tissue for their invaluable contribution to this research. All clinical data were de-identified prior to analysis to ensure patient confidentiality and adherence to ethical standards. We thank Kelly Verel, Nicole Reinholdt, Marcela White for data/sample collection. Tumor samples obtained from The Brain Tumour Tissue Bank, (London, Ontario), Cincinnati Children’s Hospital Medical Center, Children’s Memorial Hospital/Lurie Children’s Hospital of Chicago (Chicago, IL), Children’s Hospital of Los Angeles, (Los Angeles, CA).

## Authors contributions

Conceptualization R.D.,

Methodology: R.D., M.S., S.S.K.,

Formal analysis and investigation: R.D., R.S., M.S., S.S.K.

Writing-original draft preparation: R.D., K.D., R.S., R.T., B.U., R.D.

Writing, review and editing: all authors

Supervision: R.D.

---

**Supplemental Figure 1.**
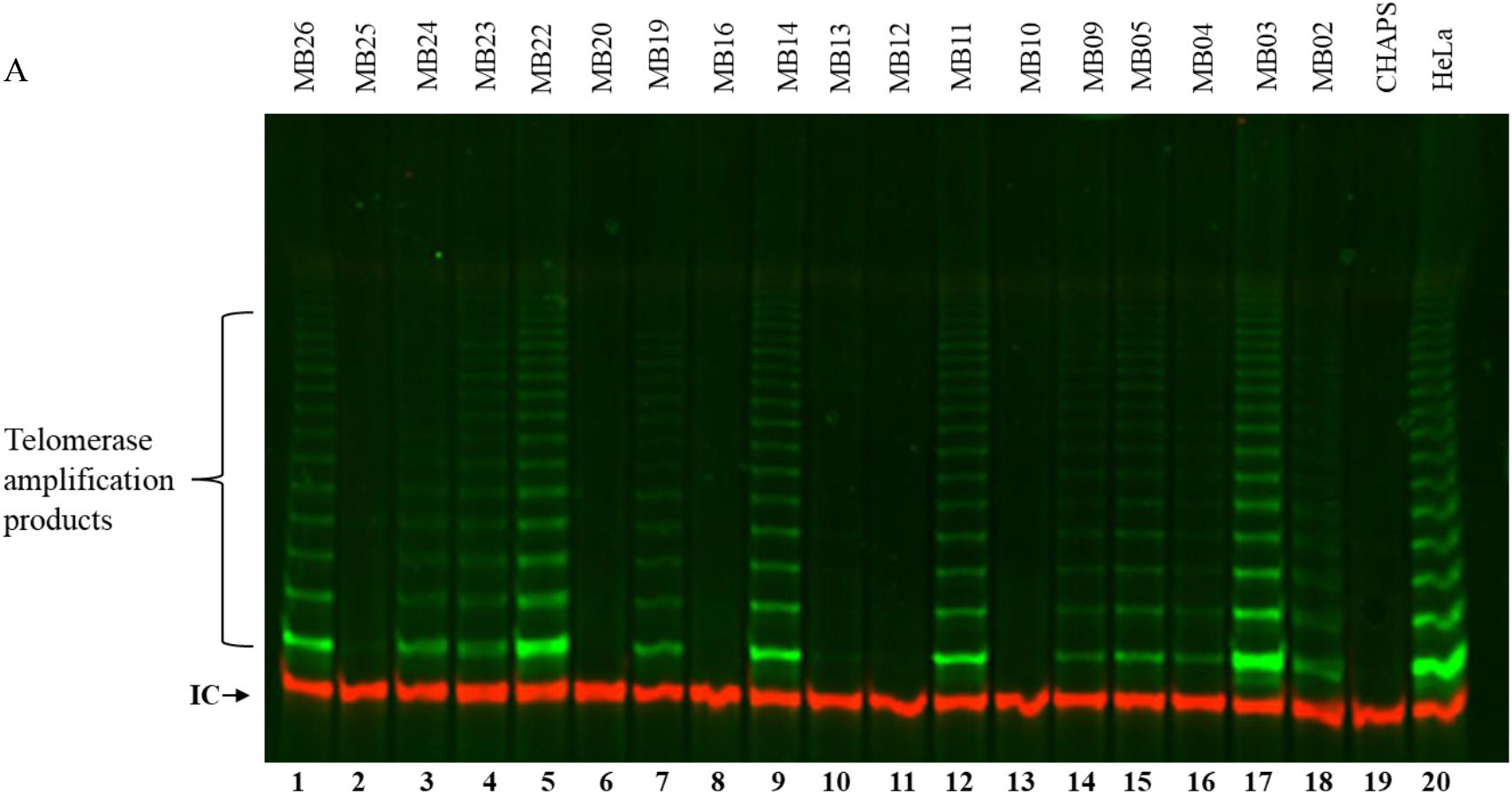
Representative TRAP assay gel image. The characteristic laddering pattern of telomeric amplification products indicates telomerase activity. HeLa cells were used as a positive control, and CHAPS lysis buffer served as a negative control. IC denotes the PCR internal control. Because the gel image is qualitative and subject to variability, telomerase enzymatic activity was quantified using fluorometric analysis and expressed as TPG units. Based on this quantitative assessment, telomerase activity was detected in 25 of 27 samples.

**Supplemental Figure 2.**
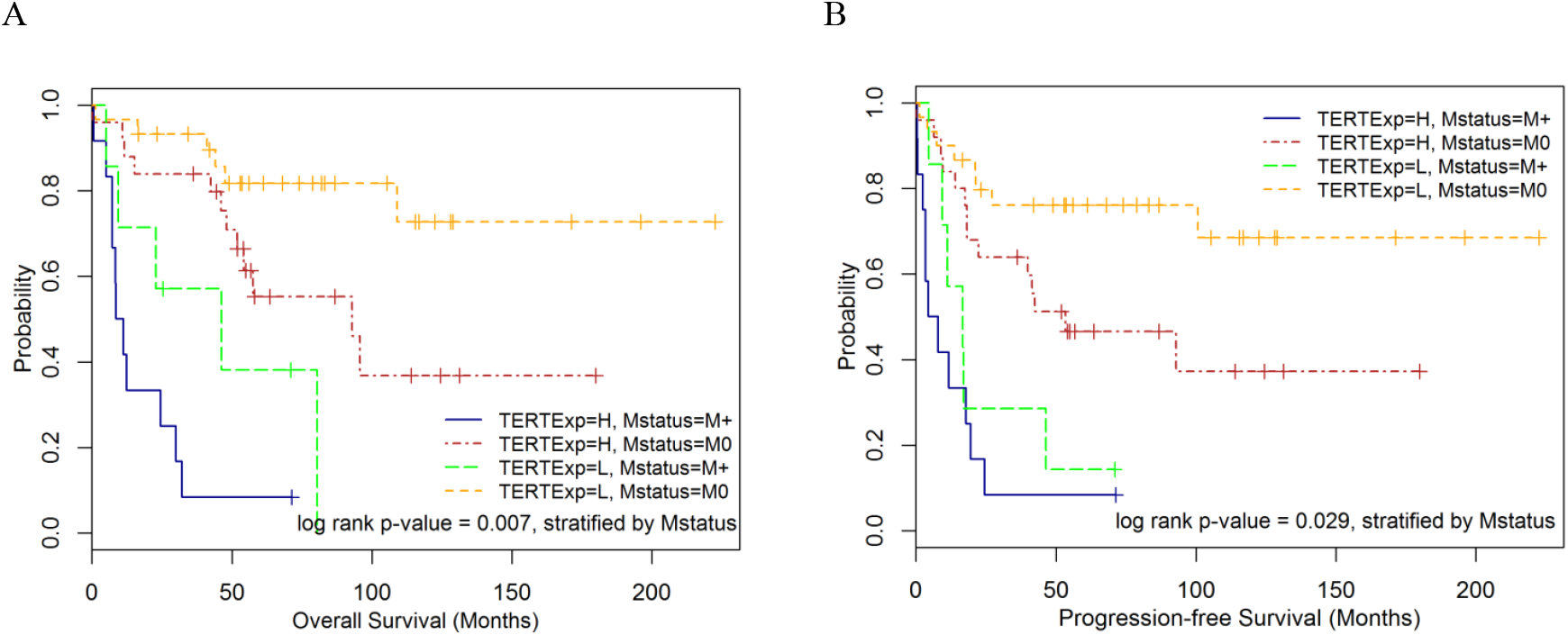
Association of *hTERT* expression and metastatic status of medulloblastoma with survival. (A-B) Kaplan-Meier plot showing the OS distribution (A) and PFS distribution (B) stratified by the *hTERT* expression and adjusting for metastatic status. Higher *TERT* expression was associated with worse OS (HR=1.35, 95% CI: 1.12-1.62, p=0.002) and PFS (HR=1.25, 95% CI: 1.07–1.45, p=0.005).

**Supplemental Figure 3.**
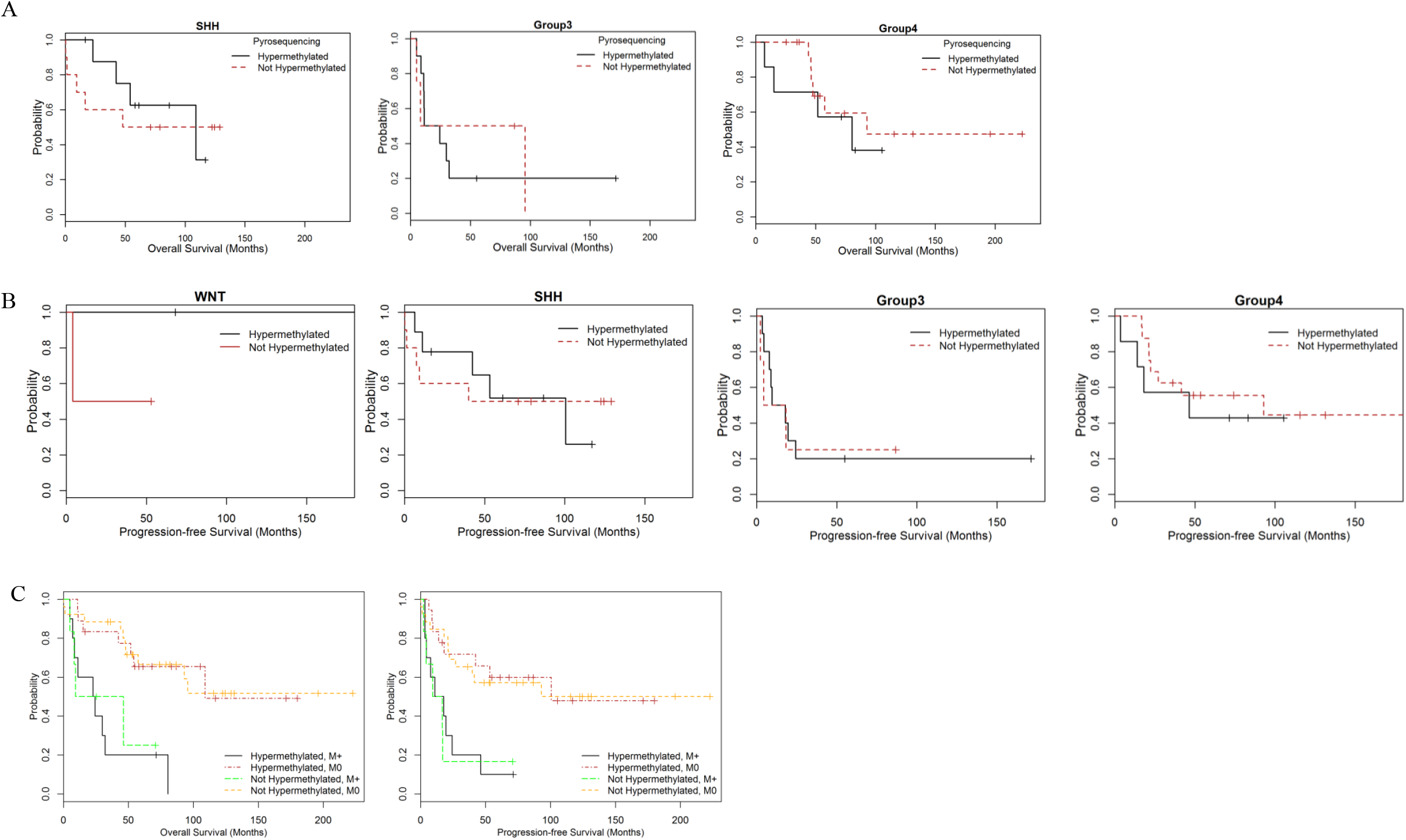
Association between *hTERT* promoter methylation status pyrosequencing and survival. All patients with well-defined subgroup (WNT, SHH, Group 3, and Group 4), non-missing OS or PFS data, and non-missing 450k array data, are included in this analysis (n = 60, 30 OS events, 34 PFS events). (A-B) Kaplan-Meier plot showing the OS distribution (A) and PFS distribution (B) stratified by the *hTERT* promoter methylation status (hypermethylation and not hypermethylated) in different medulloblastoma subgroups. (C) Kaplan-Meier plot OS and PFS adjusted for metastatic status and *hTERT* promoter methylation status

**Supplemental Figure 4.**
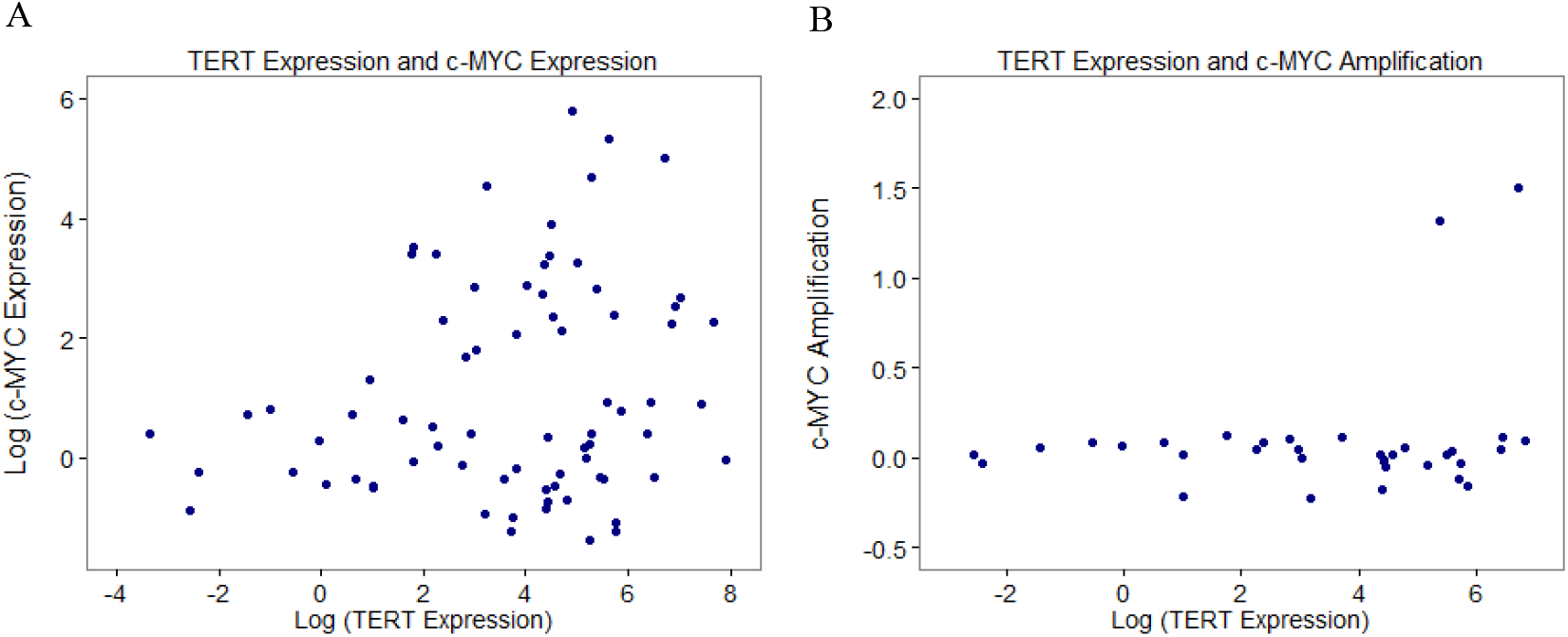
Association between *hTERT* expression and c-MYC alterations. (A) Scatter plots showing correlation between *hTERT* expression and *MYC* expression. No significant correlation was found among 74 patients with available data (Spearman’s correlation ρ = 0.15, p-value = 0.205). (B) Scatter plots showing correlation between *hTERT* expression and *MYC* amplification. Similarly, no correlation was found among 32 patients with available data (Spearman’s correlation ρ = -0.009, p-value = 0.959).

